# PacBio amplicon sequencing method to measure pilin antigenic variation frequencies of *Neisseria gonorrhoeae*

**DOI:** 10.1101/706598

**Authors:** Egon A. Ozer, Lauren L. Prister, Shaohui Yin, Billy H. Ward, Stanimir Ivanov, H Steven Seifert

## Abstract

Gene diversification is a common mechanism pathogens use to alter surface structures to aid in immune avoidance. *Neisseria gonorrhoeae* uses a gene conversion-based diversification system to alter the primary sequence of the gene encoding the major subunit of the pilus*, pilE*. Antigenic variation occurs when one of the non-expressed 19 silent copies donates part of its DNA sequence to *pilE*. We have developed a method using Pacific Biosciences amplicon sequencing and custom software to determine pilin antigenic variation frequencies. The program analyzes 37 variable regions across the strain FA1090 1-81-S2 *pilE* gene and can be modified to determine sequence variation from other starting *pilE* sequences or other diversity generation systems. Using this method, we measured pilin antigenic variation frequencies for various derivatives of strain FA1090 and showed we can also analyze pilin antigenic variation frequencies during macrophage infection.

## INTRODUCTION

Antigenic variation (Av) describes high frequency, reversible gene diversification processes resulting in the expression of many different forms of a gene product. Av is a common process in many microbial pathogens, including viruses and bacteria, and eukaryotic parasites (1–4). Gene diversification allows for stochastic population heterogeneity, which can be beneficial at the population level when selection occurs (5). While the name suggest that these systems are in response to immune surveillance, the variation can be useful for both immune and other function selection on the population level.

*Neisseria gonorrhoeae* (Gc), is a human specific organism and the sole causative agent of gonorrhea. During infection, a robust innate immune response comprised of recruited polymorphonuclear (PMN) cells and macrophages localizes to the site of infection (6,7). PMNs are the most common immune cell recruited during infection, and much of the interactions with PMNs such as recruitment and signaling have been established (8). In addition to PMNs, macrophages have been isolated from acute infection sites and Gc have been shown to modulate apoptosis and stimulate the release of cytokines and antimicrobial peptides (9–11). Gc can survive within and in the presence of macrophages, however, much remains unknown about how Gc interacts with macrophages.

To avoid adaptive immune recognition, one of the surface exposed variable proteins, the Type IV pilus, varies through conversion of the gene encoding the major pilin subunit, PilE (12,13). The Type IV pilus is required for establishing infection (14–17), so it is important that this essential factor changes throughout infection to avoid immune detection. During pilin Av, a portion of one or more donor silent copy sequences replaces part of the *pilE* gene in a non-reciprocal, homologous recombination process (18,19). There are 19 *N. gonorrhoeae* silent *pilS* copies found at various loci throughout the strain FA1090 genome (20). Any portion of the recombining silent copy, from the entire variable region to a single base can be transferred into the *pilE* locus. Recombination only requires regions of microhomology at the ends of the new sequence, and after recombination, the donor silent copy sequence remains unchanged (Figure 1). Pilin Av requires many conserved common recombination and repair factors that process gap repair and double strand breaks (21). Inactivation of some required factors, such as RecA (22), RecO, RecR and RecG, completely abrogate pilin Av (23–25), while mutation of other factors, such as RecQ, Rep, and RecJ reduce pilin Av frequencies (24,26–29).

**Figure 1.**
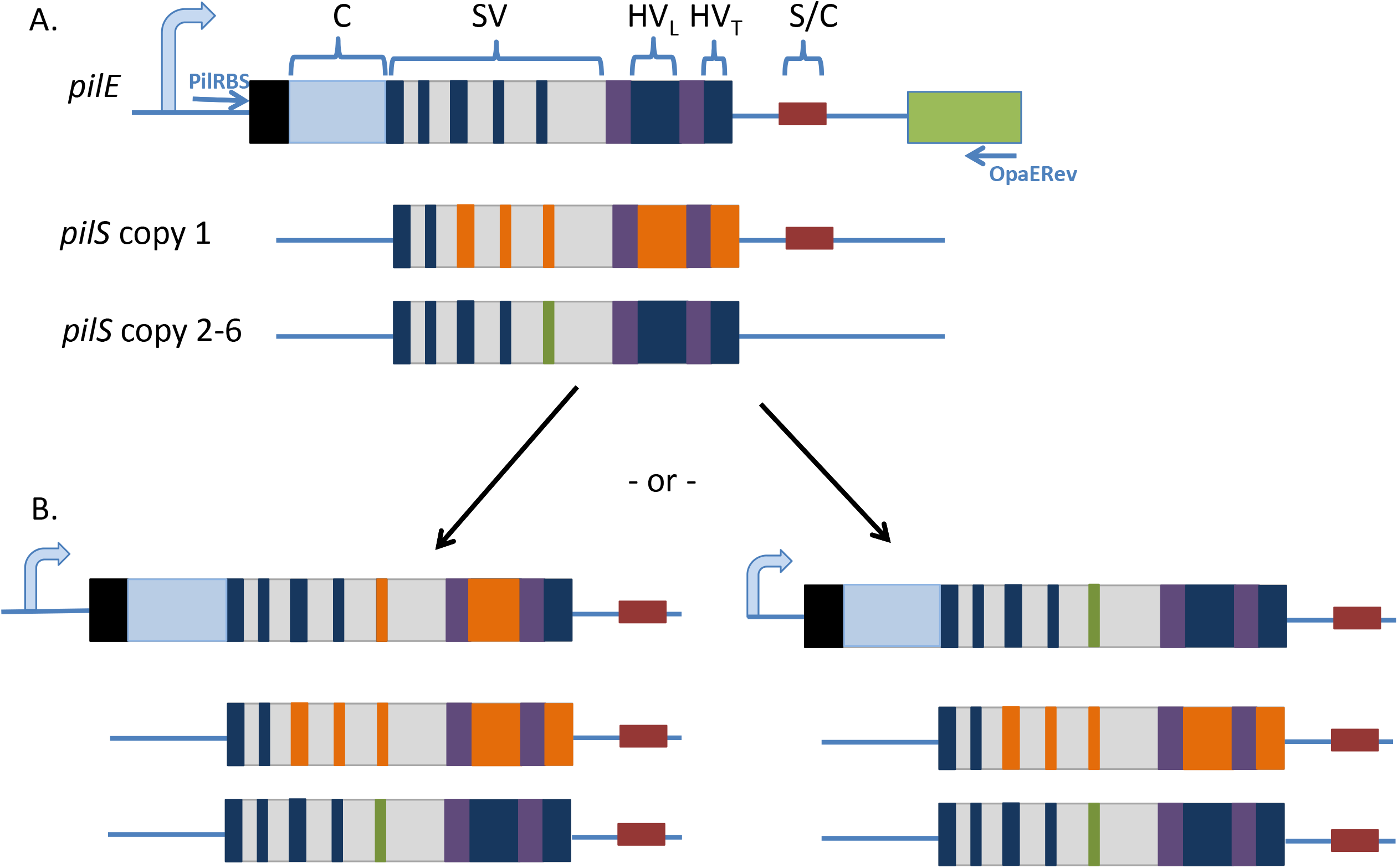
Diagram of*pilE* gene and Av process. A. Cartoon of the *pilE* gene and downstream *opa* gene (green). The 7 amino acid signal sequence necessary for proper localization of the protein (black box) is at the N-terminus of the preprotein. The constant region (C) (light blue) is not present in in any silent copy. The semivariable (SV) region has short variable regions with single amino acid substitutions (navy) in between conserved sequences (grey). The hypervariable loop (HVL) is located between the conserved*cys1* and *cys2* regions (purple) and the gene ends with the hypervariable tail (HV_T_). The 3’ untranscribed nor translated SmaCla region (S/C) (red) is found in both *pilE* and the 3’ end of all *pilS* copy 1 donors but not *pilS* copy2-6 (only one locus has six copies). The primers used for amplification, PilRBS and OpaERev are shown in blue and are not present in any silent copy. B. Gene conversion from a *pilS* copy to *pilE* can occur anywhere within the variable regions bordered by regions of microhomology. In the left example, A portion of the SV and the entire HVL sequence of the top *pilS* copy 1 donor replaces the similar *pilE* sequences but the *pliS* copy remains unchanged. In the right cartoon, the bottom *pilS* copy donates a small region of variant SV sequence.

Additionally, there are two *cis*-acting factors that function in pilin Av. An alternate DNA structure called a guanine quadruplex (G4) forms upstream of the *pilE* gene (30,31). Mutation of any of the individual GC base pairs necessary to form the *pilE* G4 structure prevents pilin Av, while mutation of any of the TA base pairs within the sequence, which are not required to form the structure, allows normal levels of pilin Av (31). Pilin Av also depends on a promoter located adjacent to the G4 forming sequence that initiates transcription within the G4 sequence resulting in a small, noncoding RNA that can only function *in cis* (32) and (Prister et al submitted).

The ~500 bp *pilE* gene contains a promoter, ribosome binding site, and conserved 5’ coding region (Constant region), which is not found in any of the silent copies (Figure 1). From bp 150-360, there is a semivariable region (SV), which contains short regions of homology interspersed with short regions of variation that are also templated in one or more silent copies. The *cys1* and *cys2* are conserved DNA sequences, which encode the disulfide bridge forming cysteines within the PilE protein (33). Located between the conserved cysteine regions is the hypervariable loop (HVL), which shows the highest amount of protein and DNA sequence diversity (Figure 1). Finally, the *pilE* hypervariable tail (HVT), from bp 477 to 492, is also highly variable in length and sequence (18). After the coding sequence, the conserved 65 bp Sma/Cla repeat is also found in *pilE* and at the end of all silent loci, and provides downstream homology for recombination (34–36).

Pilin Av leads to a change in the *pilE* sequence with anywhere from the entire variable sequence replaced to as little as one base pair altered. Pilin Av has been analyzed using a variety of methods, including: Southern Blot hybridization for the loss or gain of variable sequences, quantitative reverse transcription of specific variable sequences, enumerating the production of nonpiliated progeny, and determining the pilus-dependent colony morphology changes (PDCMC) over time (23,34,37–42). All of these methods have limitations in reproducibility and/or are affected by small changes in the growth rate. A Sanger sequencing method was developed that allowed quantification of pilin Av frequencies independent of growth rate but was limited by the low numbers of events that could be reasonably interrogated (37). The Sanger method used a Gc strain encoding an inducible *recA* allele to regulate pilin Av, and determined the pilin Av rate of 6.3 × 10^−3^ per CFU per generation for strain FA1090 *recA6*, variant 1-81-S2 (37). A Roche 454 long read sequencing method was developed to measure pilin Av frequencies that allowed large populations to be screened that also was not affected by growth rate (28). High throughput sequencing with long read lengths allows for the continuous sequencing of the entire variable region in one read. In contrast, short read technologies such as Illumina would allow determination of changes in the population, but would not be sufficient for identification of individual variants (43). Roche 454 sequencing technology is no longer available, so there is a need for new methods of pilin Av analysis with other long read sequencing technologies.

PacBio single molecule, real-time (SMRT) sequencing has been used to determine VslE Av frequencies in *Borrelia burgdorferi* (44). VslE Av is required for persistent colonization of the host because it creates a heterogeneous population that cannot be cleared by the host immune system (45). PacBio amplicon sequencing entails ligating a single stranded hairpin adaptor to each end of the amplicon, which, when denatured, creates a circular ssDNA molecule to be sequenced. However, PacBio sequencing is not very accurate with error rates around 15% per base with base mismatch and insertion/deletion errors being common (46). With circular consensus sequencing (CCS) on PacBio, each amplicon circle can be sequenced continuously to generate reads from the same template sequence several times over, leading to improvement on the accuracy of base calling. PacBio CCS was successfully used to characterize and measure *B. burgdorferi* VslE variation (47).

We developed a method to determine *Neisseria* pilin Av frequency using PacBio CCS sequencing and used this method to analyze pilin Av in strains of Gc that have previously been tested using other methods of analysis and the FA1090 common lab strain that has never been systematically analyzed before. To aid in this analysis, we developed bioinformatic methods and software to annotate variants in reads and account for the residual errors in PacBio sequencing. We report FA1090 Av frequencies in several conditions and compare our new method to previously-developed methods of analysis. Finally, we show the utility of this method by measuring pilin Av frequencies of Gc associated with human macrophages.

## METHODS

### Strains and Growth conditions

Bacterial strains used in this study were derivatives of the FA1090 clinical isolate. All gonococcal strains were screened for the human challenge isolate 1-81-S2 expressing a particular variant of the type IV pilus (48). *N. gonorrhoeae* strains were maintained on gonococcal medium base (Difco) modified with Kellogg supplements as described previously (49). FA1090 G4 mutant, *gar-10* and *gar-13* were constructed as described in Prister et al. 2019 (submitted).

### Macrophage infections

The human monocyte cell line U937 (ATCC CRL-1593.2) was cultured in RPMI 1640 media (VWR) supplemented with 10% FBS at 37C and 5% CO_2_. For macrophage differentiation, 1X10^6^ U937 monocytes were seeded per single well of a 6-well plate and were cultured in the RPMI (+10% FBS) supplemented with 10 ng/ml phorbol 12-myristate 13-acetate (PMA) for 24 hours. After that period, the media was replaced with PMA-free media and the cells were cultured subsequently for an additional 48 hours.

Differentiated macrophages were infected with the *N. gonorrhoeae* FA1090 *recA6* strain that encodes *recA* under an isopropyl-ß-D-1-thiogalactopyranoside (IPTG)-inducible promoter (50). The inoculum was prepared by selecting and streaking 4 to 5 piliated colonies as heavy patches on GCB solid media (Criterion) for 20 hours at 37°C and 5% CO_2_. Patches were collected in K+-free PBSG (PBS supplemented with 7.5mM Glucose, 0.9 mM CaCl_2_, and 0.7 mM MgCl_2_) and the number of bacteria was determined by optical density measurements.

All infections were completed in PBSG either containing or lacking potassium. In some experimental conditions, RecA expression was induced by adding 2mM IPTG (2mM) at the start of the infection and expression was validated by immunoblot analysis with a polyclonal RecA antibody (generous gift from Michael Cox, University of Wisconsin-Madison) (51). The inoculum was set at MOI of 0.2 and viable CFUs were confirmed by plating serial dilutions at the beginning (t= 0hrs) and the end (t= 12hrs) of the infection.

After the inoculum was added to the macrophage cultures, plates were centrifuged at 1000 rpm for 5 minutes to bring the bacteria in contact with the macrophages. At 12 hours post infection, ~ 4X10^8^ bacterial CFUs were recovered by directly lysing the eukaryotic cells with 0.05% Tween-20 (5 min; 37°C) and pelleting the bacteria by centrifugation (13,000 rpms for 5 min). Bacterial pellets were frozen and stored at −20°C. Equivalent number of bacteria were also prepared from the inoculum (~5X10^8^ CFUs) and frozen. For *pilE* Av frequency analysis, genomic DNA was isolated from the frozen pellets with GenElute Bacterial Genomic DNA Kit (Sigma-Aldrich).

### Preparing strains for PCR amplification and sequencing

Strains were struck out from frozen stocks and grown overnight for 18 hours. A single colony was picked using a sterile 6 mm filter disk and dispersed in 500 μl GCBL by vortexing. The isoated colony was diluted in GCB medium and different dilutions plated on GCB solid medium to obtain 200-400 colonies per plate. The remainder of the cell suspension was pelleted, washed with 1X PBS and bacteria lysed in cell lysis buffer. The lysed bacteria were then used as a template for PCR and subsequent sequencing with primers PilRBS and Sp3A. This step ensured that the strains all started as the same variant *pilE* sequence (1-81-S2) (48).

The colonies were grown for 22 hours and number of colonies recorded. At least 1000 colonies (from 1041-1255 colonies) were pooled in GCBL and genomic DNA was isolated using Qiagen QiaAmp kits. The genomic DNA was used as a template for PCR and subsequent sequencing with primers PilRBS and Sp3A to determine whether the majority starting sequence was retained (48). Genomic DNA was amplified using the following reaction: 1 ng genomic DNA, 20 μM dNTPs, 1X Phusion reaction buffer, 0.5 μM Primer 1, 0.5 μM Primer 2, 3% DMSO, 1 unit Phusion Hot Start Flex (NEB) Polymerase and ddH_2_0 (Supplemental Table 1). The reaction was run under the following conditions: 98° C for 30 seconds for initial denaturation and polymerase activation, 98° C for 10 seconds, 65° C for 30 seconds then 0.3° C reduced in each cycle, 72° C for 1 minute, repeat cycles 30 times, final extension for 5 minutes. For each sample, a different 16 base barcode was used on both the forward and reverse primers, PilRBS-TTTCCCCTTTCAATTAGGAG and OpaERev-GGGTTCCGGGCGGTGTTTC leading to a 788 bp product, 820 bp with barcodes (Supplemental Table 1).

The PCR products were run on an agarose gel without ethidium bromide and UV exposure. Gel extraction was performed with the QiaQuick Qiagen gel extraction kit, but with the gel slices dissolved at room temperature to maintain DNA integrity. The columns were eluted with TE buffer and pooled to obtain 300 ng of DNA per sample (Figure 2). Samples were then submitted to University of Maryland Genomics resource center where the amplicons were purified with SPRI clean up, quantified and combined into two pools. SMRTbell library prep was performed and the pools were sequenced on Pacific Biology Sequel SMRT cells with v3 reagents.

**Figure 2.**
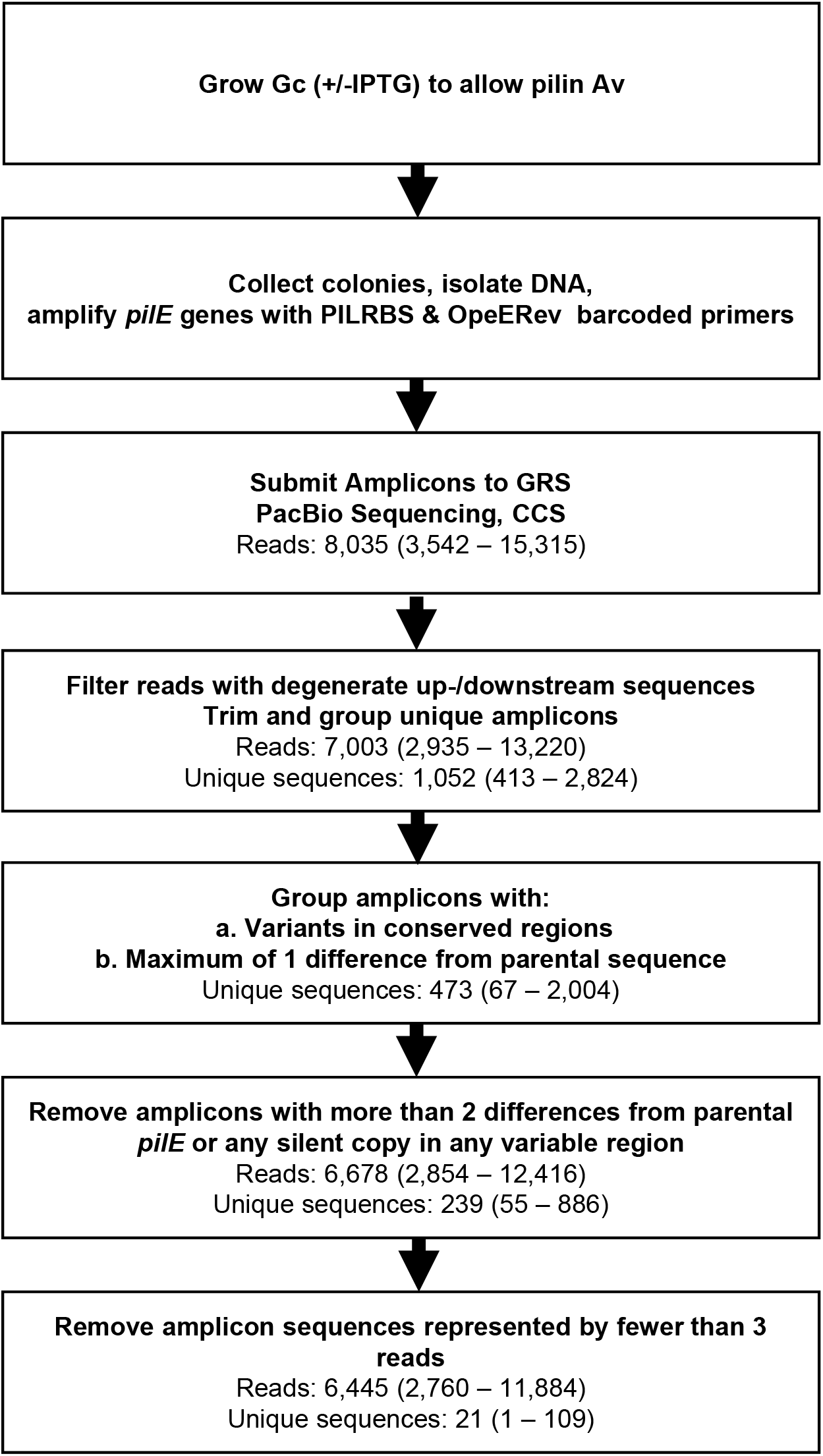
Flow chart of protocol and results from amplicon read processing. Strains were grown for 22 hours and colonies pooled to isolate genomic DNA. The *pilE* gene was amplified with primers containing a specific barcode for each condition or sample. The PCR was gel purified and sent to the Genomics Resource Center for library preparation. The reads were then demultiplexed and analyzed using SwitchAmp as described in the methods. The flow chart describes the filtering steps used for amplicon read processing in SwitchAmp. The average number of reads in each step is displayed below each step with the range in parentheses for each pool.

### Processing Reads and Aligning Pilin Variants

PacBio subreads generated for each amplicon were converted to Circular Consensus (CCS) reads using ccs v3.0.0 and demultiplexed using the SMRT Analysis software module (Pacific Biosciences, San Francisco). CCS reads for each amplicon pool were then filtered and analyzed using the software package SwitchAmp reported here. The SwitchAmp program is written in Perl and C and can be downloaded from https://github.com/egonozer/switchAmp. The program is run from the command line and compatible with Macintosh and Linux operating systems. Inputs are the fastq-formatted PacBio CCS read file of amplicon sequences for an experiment and a file listing the positions of variable regions relative to the parental sequence and the sequences of the silent copies at these positions (Supplemental Table 2). This variable region file can be generated manually or from an alignment of the parental sequence to the silent sequences using the script fasta_alignment_conserved.pl that is provided with SwitchAmp. Briefly, SwitchAmp performs the following steps: 1) Filters and removes flanking sequences in each read surrounding the *pilE* gene sequence. 2) Orients amplicon sequences and groups reads with identical sequences. 3) Aligns read sequences to the parental sequence using the Needleman-Wunsch algorithm. 4) Parses variable regions in aligned read sequences to identify matches against the provided parental or silent sequences for each variable region. 5) If a variable region does not perfectly match a provided parental or silent sequence, parental or silent copy sequences with the smallest Levenshtein distance to the read sequence will be determined. 6) If the total distance from the parental sequence across all variable regions in a read is less than or equal to a given cutoff, all reads with this sequence will counted as parental. 7) Pairwise Levenshtein distances are calculated for concatenated sequences of all variable regions in each unique read sequence and hierarchical clustering is performed (Figure 2). The outputs of the program include a table with each unique read sequence, its total frequency in the input read file, and the closest matching silent copies in each variable region. The table also shows the Levenshtein distances between each read sequence and the closest matching silent copy or copies in each variable region. Other program outputs include a file with all unique read sequences, a fasta-formatted file for each variable region listing all variable sequences identified, a newick-formatted tree file with the dendrogram generated by hierarchical clustering of all variable region sequences, and a file summarizing the results of each filtering and processing step by the program.

## RESULTS AND DISCUSSION

### Optimization of conditions for PacBio sequencing

To enable measuring of pilin Av frequencies by PacBio sequencing, strains were grown for 22 hours on GCB plates, which results in ~19-20 generations (28). The IPTG-inducible *recA6* strains were grown on solid medium with IPTG to allow for pilin Av. All of the strains tested had similar growth rates, so all Gc were harvested at the same time point (28). Between 1041-1255 colonies were pooled, and genomic DNA was extracted (Figure 2).

False “variants” can be produced by in vitro recombination during PCR amplification when an extension intermediate primes a different silent copy extension intermediate producing a hybrid sequence of parent and silent copy (28,52). This has also been seen in other systems such as analysis of drug resistant allele variants of HIV (53). To limit these PCR artifacts, low genomic template (1 ng), a high processivity polymerase (Phusion Hot Start Flex), and the touchdown PCR cycles were used. Touchdown PCR initiates with high annealing cycle temperatures, which are slowly lowered each annealing cycle to ensure specific primer binding of the correct region. Using all these techniques, PCR recombination was reduced to background levels (Table 1). PacBio barcodes (Supplemental Table 1) were used to differentiate the samples, and to minimize PCR cycles, the barcodes were added to the primers for the *pilE* gene, OpeERev and PilRBS (48,54). Each pairs of barcoded primers was tested and any pairs showing aberrant products were not used. After PCR, the products were gel purified from gels without ethidium bromide by staining a reference lane to localize the product and DNA was extracted with the QiaQuick gel extraction kit without heating the gel slices. The University of Maryland Genomic Resource Center conducted the further steps, including PCR purification using solid phase reversible immobilization and quantification by Qubit. The pooled samples were prepared for sequencing with the SMRTbell library kit from PacBio and run on the Sequel system (Figure 2). After sequencing, reads were demultiplexed and consensus sequence was determined by SMRT Analysis Software with a read confidence of 90, minimum read of three passes and minimum read length of 50. The average amount of total sequence generated per condition was 6.32 Mbp (range 2.76 – 11.91 Mbp). The average read length was 786.7 bp with a median read length of 789 bp. In all samples, including those from total macrophage/Gc DNA there was sufficient number of reads per sample to have a measure of pilin Av frequencies. Read characteristics and NCBI accession numbers are shown in Supplemental Table 3.

**Table 1.** Pilin Av frequencies of strain FA1090*recA6*, 1-81-S2. Pilin Av frequencies from PacBio Sequencing and SwitchAmp analysis as compared to those reported by Rotman et al. (28) using 454 method and a different computational program. The pilin Av frequency was calculated by dividing the number of *pilE* variant reads by total *pilE* reads after filtering for quality. The number of reads listed is the final reads after all filtering steps listed in Figure 2. ND – not determined

|  | **Pool A** |  | **Pool B** |  |  | **Pool A** |  | **Pool B** |  |
| --- | --- | --- | --- | --- | --- | --- | --- | --- | --- |
| **PacBio** | **Reads** | **% Av** | **Reads** | **% Av** | **454 \*** | **Reads** | **% Av** | **Reads** | **% Av** |
| ***recA6* - IPTG** | 8881 | 0.17 | ND | ND | **Av deficient** | 96,993 | 0 | ND | ND |
| ***recA6* + IPTG** | 4001 | 4.65 | 11884 | 6.70 | ***recA6* + IPTG** | 6,495 | 10.6 | 5,977 | 10.8 |

### SwitchAmp Program for Variant Analysis

To analyze the sequencing reads, we identified all regions that can differ between the 1-81-S2 *pilE* sequence and all the possible changes that could occur from any of the 19 silent copies (Supplemental Table 2), since all variation events should be templated from a silent copy. For all samples, the average number of CCS reads generated per amplicon set was 8035 reads (range 3542 - 15315) (Figure 2 and Supplement Table 3). On average, 13% of reads (range 9.8% − 20.4%) were filtered for exceeding a maximum of two mismatches in either the upstream and/or the downstream sequences flanking the *pilE* gene sequence. This analysis resulted in an average of 7003 amplicon sequences in total per sample (range 2935 – 13220). Amplicon sets were filtered to group all amplicons with either isolated conserved region mismatches or fewer than two base mismatches across variable regions with the parental sequence. Amplicon sets were then further filtered to remove sequences that had more than two base differences from parental sequence or any silent copy. This filtering resulted in an average of 6678 sequences per read set (range 2854 − 12416). We chose to remove amplicon sequences represented by fewer than three reads in each set to reduce potential error from low frequency variants resulting from sequencing errors. Using this cutoff resulted in removal of an average of 93.8% of unique sequences (range 87.2% − 98.9%) in each read set, however those unique sequences only represented 3.5% of all previously-filtered reads (range 1.0% − 13.2%). These results indicate that the SwitchAmp algorithm effectively identifies and characterizes high quality Av sequences from long read amplicon sequencing experiments.

### Pilin Av frequencies in IPTG-regulated *recA6* strains

Strains with the IPTG regulatable *recA6* allele were grown for 22 hours as described previously (28). Without RecA induction, we measured 0.17% pilin Av (Table 1). We have repeatedly analyzed pilin Av in this strain without IPTG induction and have never recorded a true pilin Av event over many studies. Therefore, we conclude that these variant reads are the result of sequencing errors being recorded as true events or alternatively the result of PCR recombination. The pilin Av frequency with IPTG induction of RecA was 4.65% and 6.70% in two biological replicates. The 2% difference in the frequency of pilin Av measured between the two biological replicates is most likely due to the stochastic nature of pilin Av and the fact that events that occur early will be overrepresented in the population. These data highlight the importance of biological replicates and demonstrates that PacBio can be used to measure pilin Av in Gc. These pilin Av frequencies are lower than reported using 454 sequencing and a different analysis method (Table 1). The 454 sequencing had a different error rate (0.49% per base) (55) and in the absence of IPTG had a higher background rate of about 1%. With different error rates, we may be discounting actual variants if there are errors in other regions of the *pilE* gene and the sequence is then discarded. The two methods also used different computational analysis methods, which calls variants differently.

### Pilin Av frequencies in an unregulated FA1090 strain

All previous pilin Av sequencing studies have used the *recA6* strain to start with a uniform population and to limit Av to a specific number of generations. Measuring pilin Av frequencies of FA1090 with an unregulated *recA* gene has never been reported. The difficulty with measuring Av in FA1090 without the *recA6* allele is that *pilE* is constantly varying during growth and the experiment cannot start with a single variant. Therefore, it is likely that early Av events will occur and predominate in the population. We used this PacBio method to measure Av in FA1090. Gc strains were grown overnight on solid medium from freezer stocks for 18 hours. Several single progenitor colonies were each plated onto solid medium and grown for 22 hours. In parallel, the *pilE* of each progenitor colony was sequenced by Sanger sequencing and only progenitors with the *pilE* variant 1-81-S2 were processed for PacBio sequencing. Sanger sequencing only gives a population level sequence of the most common base at each position and since Av frequencies are approximately 10% across the variable regions of a gene (34), this low frequency variation at a population level cannot be detected by standard sequencing and we are certain that there was always a population of variants that arose during the growth of the progenitor colony.

As anticipated, the continual pilin Av frequencies measured for FA1090 were higher than those of *recA6* strains (Table 2). The pilin Av frequency of the two FA1090 biological replicates were 17.90% and 17.40%. Mutation of the G4 forming sequence or the promoter of *pilE* G4 sRNA (*gar*) are required for Av (31,32) produced pilin Av frequencies of 0.21% and 0.1%, respectively (Table 2), which is similar to the pilin Av frequency of the *recA6* strain without *recA* induction (Table 1). Mutation of the −35 sequence of the G4 sRNA promoter (*garP-_35_*) in the same FA1090 background showed reduced levels of pilin Av of 6.13 and 5.73%, which is consistent with the reduced levels of *gar* RNA produced when the −35 sequences was mutated. (Table 2) (Prister et al. 2019 submitted). We assume that these levels of pilin Av represent steady state levels under these growth conditions.

**Table 2.** Pilin Av frequencies of FA1090 strains. Av frequency of the FA1090 strain were calculated similarly to Table 1 using PacBio sequencing and analysis with SwitchAmp. ND – not determined

|  | Pool A |  | Pool B |  |
| --- | --- | --- | --- | --- |
|  | Reads | % Av | Reads | % Av |
| **FA1090** | 7296 | 17.61 | 5396 | 16.9 |
| **G4 mutant** | 7252 | 0.21 | ND | ND |
| **FA1090 *garP*<sub>-10</sub>** | 4707 | 0.1 | ND | ND |
| **FA1090 *garP*<sub>-35</sub>** | 4522 | 6.13 | 4434 | 5.73 |

**Table 3.** Analysis of potential P-sequences. We analyzed the number of variants that contained P-sequences, and therefore would produce an under-piliated colony and could outgrow during the 22 hour growth. The number of P-variants was divided by total variants to obtain the %P-. ND – not determined

|  | Pool A |  | Pool B |  |
| --- | --- | --- | --- | --- |
|  | % Av | %P- | % Av | %P- |
| ***recA6* 0 IPTG** | 0.17 | 0 | ND | ND |
| ***recA6* 1 mM IPTG** | 4.65 | 12 | 6.77 | 32.8 |
| **FA1090** | 17.61 | 41.6 | 16.9 | 26.8 |
| **G4 mutant** | 0.21 | 20 | ND | ND |
| **FA1090 *garP*<sub>-10</sub>** | 0.1 | 0 | ND | ND |
| **FA1090 *garP*<sub>-35</sub>** | 6.13 | 29 | 5.73 | 28 |

These results demonstrate that PacBio amplicon sequencing can measure differential pilin Av frequencies in FA1090 strains without the use of the inducible *recA* construct. Both biological replicates of FA1090 were very similar in Av frequency, but the *recA6* strains did have some variability between replicates, highlighting the need for replicates. Since FA1090 populations can undergo Av at any time point during the experiment, there is potential for a “jackpot” event to occur very early in growth and result in a large portion of the bacteria containing the variant sequence. Therefore, it is necessary to perform biological replicates to verify that Av frequency measurements are consistent. This method may not be able to differentiate small differences, because there is still variation between biological replicates.

### Analysis of silent copy donors

We determined which silent copies were used during pilin Av, and if any of the mutants analyzed had different patterns of donor silent copy usage. Previously, no mutation has been shown to alter silent copy choice (28,37). The SwitchAmp software we developed identifies the most common silent copy sequence among all the regions of variation in each amplicon sequence. This analysis can allow for the identification of the donor silent copy in a variant, because if one region has a sequence change common among many silent copies, and the next region also has a change that matches a single silent copy, the most likely donor copy can be inferred. For example, if in one read, one variable region has a sequence that is identical in silent copies 1c1, 1c2,1c3, and 1c4 and the next downstream variable region has a sequence that is found only in silent copy 1c2, then the most likely donor was 1c2. SwitchAmp also provides a table of the silent copy choice for each variable region in each variant sequence, so a more in-depth analysis can be performed with manual inspection.

There were four variants most commonly seen among our parental strains (Figure 3). The most common variant in most strains was the change to the identical *pilS2* copy 1 (2c1) or *pilS6* copy 1 (6c1) in variable region var37, which is also called the hypervariable tail (HV_T_) (56). The next most frequent donor was *pilS3* copy 1 (3c1), mostly in region var32, which is also called the hypervariable loop (HV_L_) These donors are the same as the most frequent report previously using this same strain and starting *pilE* sequence (12,18,28,37,57). Another category of variants was multidonor, which refers to sequence changes common to multiple silent copies. There are many regions in the silent copies that have shared sequences, therefore the exact donor sequence cannot be determined. Additionally, a double recombination event could also be part of this population when both recombination regions have a similar length, for example if region 3 and 4 have 3c1 sequence and regions 15 and 16 have sequence from 1c1, the output would include both sequences, and be termed multidonor. However, for example, if 3c1 was the donor for regions 3, 4, and 5, and 1c1 was the donor for 15 and 16, the output would only include 3c1 because it was the most common silent copy used across the whole sequence.

**Figure 3.**
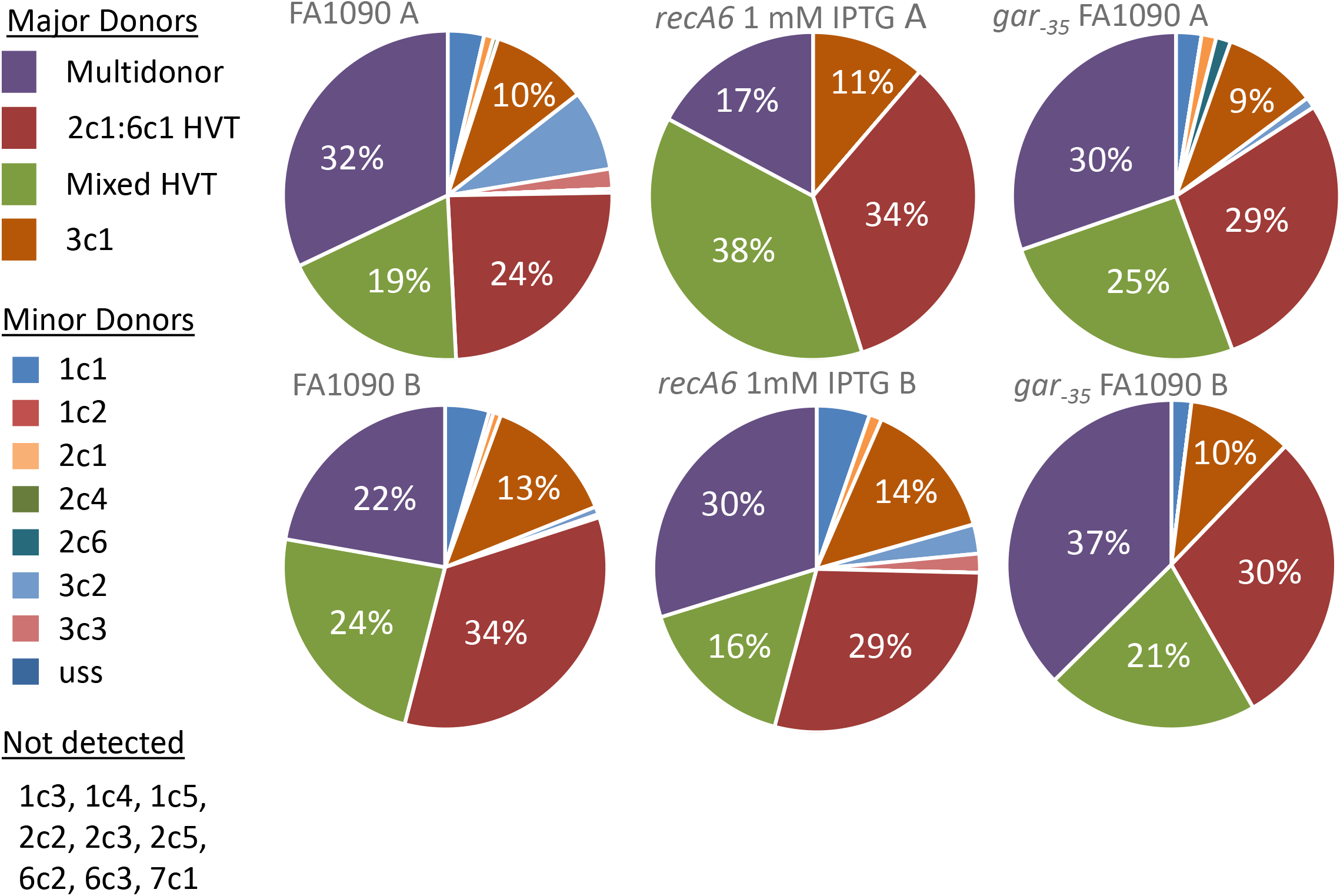
Donor silent copies. The most common silent copy found in each variant sequence was determined for strains FA1090, *recA6* 1 mM IPTG, and FA1090 *gar_-35_,* Multivariant indicates that the changed sequence was common to multiple silent copies, and there was no dominant silent copy. HV_T_ is the hypervariable tail or var37 in our analysis. This is the most common region of variation. Mixed HV_T_ indicates that the tail contained a mosaic sequence. The minor silent copies were used at a much lower rate and some silent copies were not used in our analysis.

As found with 2c1/6c1 variants discussed above, other recombination events also involved the HV_T_. Many HV_T_ variant sequences were mosaic sequences, containing sequences that mostly matched the parental 1-81-S2 sequence, but also had a few nucleotides that would have been donated from 2c1/6c1. These mosaics were common in strains undergoing pilin Av but were never seen in the Av deficient control samples. We propose that these mosaic sequences are the result of one strand of a silent copy annealing with the 1-81-S2 *pilE* tail during the process of pilin Av. Alternatively, they could be formed as regions of heteroduplex when a Holliday junction is formed. If one strand of DNA is from 1-81-S2 and the other from a silent copy, there are regions upstream and downstream of the HV_T_ with homology and throughout the HV_T_, the duplex will contain mismatches. The mismatch correction system could then correct the mismatches to either parent or silent variant sequence creating a mosaic tail. We have previously shown that the RuvABC enzyme that processes Holiday junctions is necessary for pilin Av (58) and that mismatch correction controls the frequency of pilin Av (59).

Currently, SwitchAmp does not precisely identify recombination events where two different silent copies donate sequence at different locations in the gene, although the program output can be manually examined to find amplicon sequences representing double recombinants. Additionally, as there are many regions of microhomology among silent copies, if there is a recombination event in one of these regions, the program can detect the variation, but not assign a specific donor copy. These regions will output as a list of all the possible donors in each variable region and are tallied as multidonor in the table.

### Nonpiliated (P-) colony morphology affects pilin Av frequencies

Pilin Av events can result in a colony morphology change when a premature stop codon is incorporated into the coding sequence. Additionally, some combination of silent sequences can produce a non-productive PilE protein that cannot efficiently assemble into the pilus fiber or create a stable pilus fiber leading to a nonpiliated colony phenotype (60). Either of these events can alter the colony morphology, and more importantly, increase the growth rate of Gc (60,61). In order to better understand whether pilin Av events leading to a nonpiliated (P-) phenotype were over-represented in some samples and could possibly explain some of the Av frequencies, we first identified P- associated sequences based on previous studies (Supplemental Figure 4) (37,62). We used SwitchAmp to identify variants containing P- associated sequences and determined the number of potential P- variants as a factor of the total Av variants in each strain (Table 3).

Overall, there was a slight increase in the proportion of P- sequences in strains where pilin Av frequencies were increased, but not enough to conclude that P- sequences are strongly favored when pilin Av frequencies are higher. One would expect that if there is no growth benefit selection for non-piliated Gc, the proportion of P- variants in the total variant population would be similar regardless of Av frequency. For example a P-frequency of 20% when the pilin Av frequency is 4% and 20% P- when the frequency is 14%. In the *recA6* strains in which recombination was induced with IPTG, we saw an increase in pilin Av frequencies between biological replicates (20%). The higher pilin Av frequency correlated with a much higher proportion of P- sequence, which may explain some of the differences in frequencies (Table 3). We would predict that when P- variants arise early during IPTG induction, the proportion of P- variants will be higher and their growth advantage will amplify their representation. This result contrasts with the P-% in the FA1090 biological replicates. The two FA1090 replicates have very similar pilin Av frequencies, but there is a 15% different in P-% between the replicates. We would hypothesize that these P- sequence changes occurred late in the 22 hour timeframe, which does not allow the high P-population to outgrow before we collected the DNA. Since our analysis pooled many colonies, and we cannot detect P- variant within a colony, we cannot independently test these ideas. The Sanger sequencing method of pilin Av previously used was able to report the colony morphology for each variant, which is one benefit to that method (37).

### Measuring pilin Av in association with human macrophages

One of the advantages of this method of measuring pilin Av in a population of bacteria is the ability to do an analysis within a complex biological context. We tested whether we could measure the frequency of pilin Av during infection of macrophages, a cell type encountered during infection (9). Pilin Av has been detected from isolates within the human host (48,63). Reduced iron availability has been shown to increase Av frequencies (64), but no other external signal has been found to influence pilin Av rates. Pilin Av frequencies do not change when Gc infect T84 epithelial cells (65), however, pilin Av has never been measured in the context of macrophage infection.

The FA1090 1-81-S2 *recA6* strain was added to differentiated U937 macrophages at an MOI of 0.2 with IPTG in the tissue culture medium and total DNA was isolated after 12 hours of infection. IPTG induction of *recA* was confirmed by immunoblot analysis using anti-RecA antisera (Figure 4). Potassium was also added to the media to dampen the macrophage inflammasome response. The *pilE* gene was amplified from the total DNA after 12 hours of association. Because the inoculum was prepared in the absence of IPTG, as expected Av was not observed when RecA expression was not induced. Conversely, IPTG-treated Gc-macrophage cocultures produced pilin Av frequencies of 1.2% and 1.68% in two biological replicates (Table 4). When potassium was added in the media to dampen the macrophage inflammasome response the Av frequencies were slightly reduced to 1.1% and 0.69% respectively (Table 4). The spectrum of silent copy donors in macrophage infections and in vitro cultures were similar (Figure 5 compared to Figure 3). However, due to the small number of variation events (between 19 and 125), the proportion of each event is different than those measured in monoculture (Figure 5). Based on this initial analysis, Gc do vary in the presence of macrophages and silent copy choice is similar to plate grown bacteria indicating that the silent copies used during plate mirror those that occur during infection.

**Figure 4.**
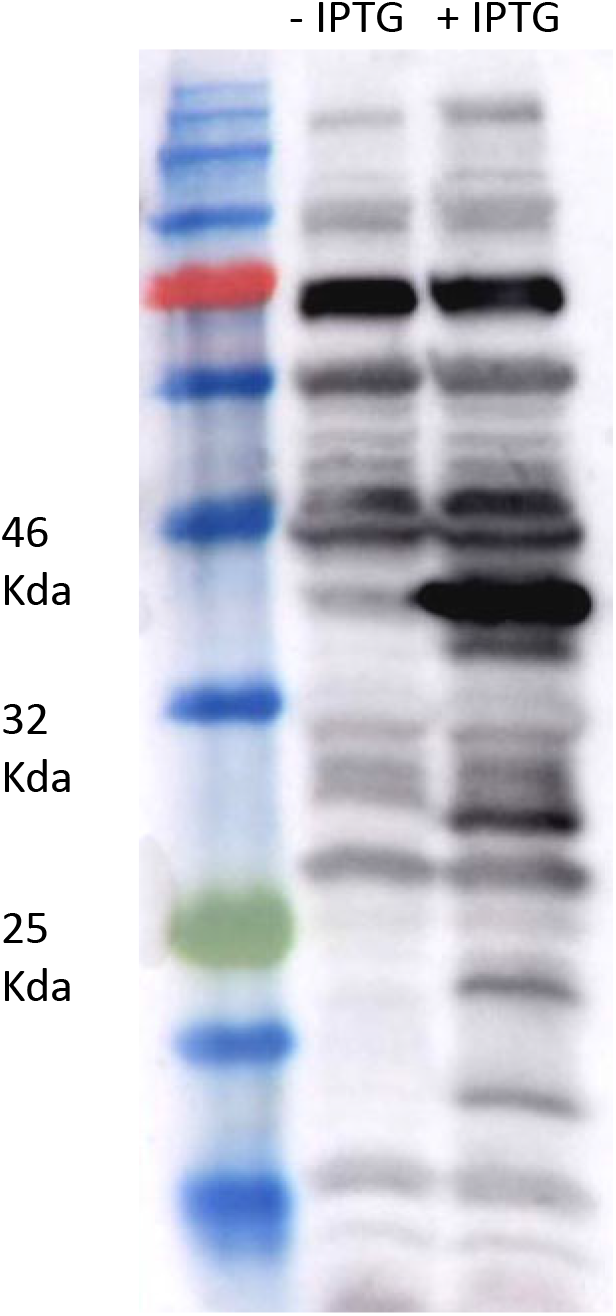
Analysis RecA induction during macrophage infection. Western blot using a polyclonal RecA antibody. Upon addition of IPTG, the RecA band at 40 Kda appears. Based on previous results, the faint band at 40 kd in the –IPTG lane is another protein. Ladder labeled for reference.

**Table 4.** Pilin Av frequencies during macrophage infection. The Av frequencies (Av %) were calculated by dividing the number of variant reads by the total reads after filtering by the SwitchAmp program. All varying strains were analyzed in biological replicates (Repeat A and B).

|  | **Pool A** |  | **Pool B** |  |
| --- | --- | --- | --- | --- |
|  | **% Av** | **Reads** | **% Av** | **Reads** |
| **Inoculum** | 0.0 | 6405 | 0.1 | 7972 |
| ***reca6*, no IPTG** | 0.0 | 7350 | 0 | 5367 |
| ***reca6*, 2mM IPTG** | 1.2 | 5733 | 1.68 | 7437 |
| ***reca6*, +K, no IPTG** | 0.0 | 5049 | 0.0 | 7614 |
| ***reca6*, +K, 2mM IPTG** | 1.1 | 8385 | 0.69 | 2760 |

**Figure 5.**
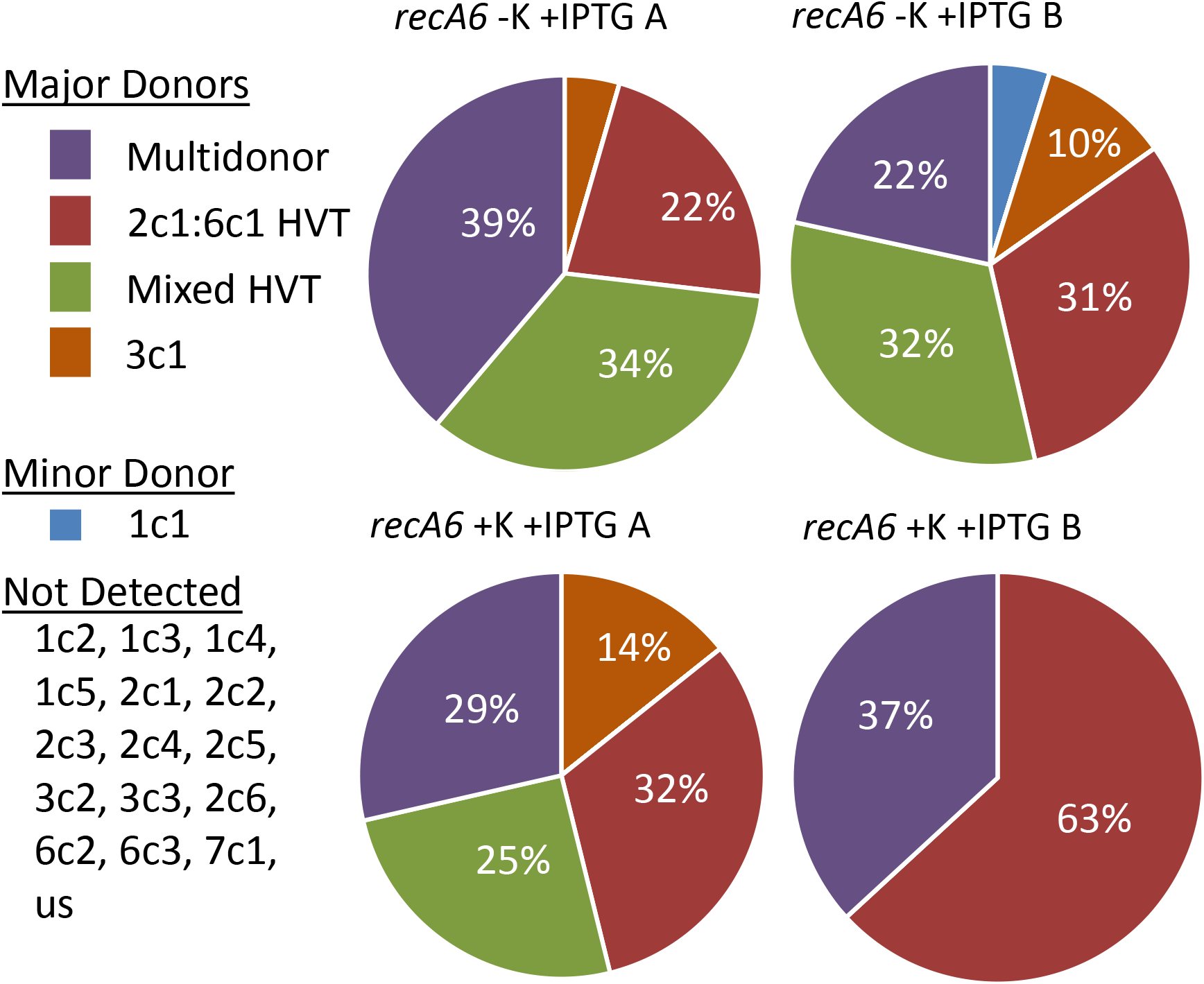
Silent copy choice during macrophage pilin Av. Each pie graph represents all variation events for each of the samples that contained pilin variants during macrophage infection. There are four major donor silent copies in most strains, the HV_T_ change to 2c1 or 6c1, where the exact donor cannot be determined because both donors are identical in that region. The creation of a mosaic tail sequence also originated from 2c1 or 6c1. Multidonor encompasses variants that have changes that match multiple silent copies, or there was no single major donor such as a double recombination event. There were only 19-125 variation events recorded in each sample.

This is the first detection of pilin Av during macrophage infection and proves this method can work as a measure of pilin Av when colony morphology cannot be observed. Both biological replicates are similar at this 12 hour timepoint. In the future, we can use this method to determine if the frequency changes during infection, but this would require the number of generations to match from plate grown to infected bacteria and account for some bacterial death during infection. The addition of potassium did slightly lower the Av frequencies, however more experiments are needed to determine whether there is a true effect of potassium addition on Av frequencies and whether inflammasome activation could influence the frequency. Regardless, this analysis is the first to show Av does occur in macrophage associated Gc and paves the way for future experiments investigating pilin Av in the context of infection.

### Conclusions

PacBio, long read amplicon sequencing is an effective means to analyze Gc pilin Av frequencies. This method will be most useful for strains that grow at different rates or when growth rates cannot be accurately calculated, such as during infection. In contrast to short read sequencing approaches, this method reports the spectrum of silent copy donor in addition to the total number of variation events. To make this method adaptable to other scenarios, we created the SwitchAmp program to report pilin Av frequencies. As discussed in the introduction, there are many systems that undergo gene diversification, such as *Borrelia*, Trypanosomes, or even antibody generation. PacBio sequencing with the SwitchAmp program allows for the user to input all variable regions and possible sequence changes and the program then analyzes all amplicon reads, so it could be used to analyze many different gene diversification systems.

## AVAILABILITY

SwitchAmp and associated software can be found at https://github.com/egonozer/switchAmp with documentation.

The amplicon sequencing data is available through the NCBI sequencing read archive (SRA), https://www.ncbi.nlm.nih.gov/sra, under SRA study accession number SRP214219. Accession numbers for individual read sets are given in Supplemental Table 3.

## SUPPLEMENTARY DATA

Supplementary Data are available at NAR online.

## ACKNOWLEDGEMENT

We thank the Northwestern University Center for Genomic Medicine, NUSeq Facility for processing the Sanger sequencing samples and University of Maryland Genomic Resources Center for PacBio sequencing and library preparations.

